# Mixture Models for Domain-Adaptive Brain Decoding

**DOI:** 10.1101/2025.10.05.680511

**Authors:** Aidan Dempster, Brokoslaw Laschowski

## Abstract

A grand challenge in brain decoding is to develop algorithms that generalize across multiple subjects and tasks. Here, we developed a new computational framework to minimize negative transfer for domain-adaptive brain decoding by reframing source selection as a mixture model parameter estimation problem, allowing each source subject to contribute through a continuous mixture weight rather than being outright included or excluded. To compute these weights, we developed a novel convex optimization algorithm based on the Generalized Method of Moments. By using model performance metrics as the generalized moment functions in our GMM optimization, our algorithm also provides theoretical guarantees that the mixture weights are an optimal approximation of the importance weights that underlie domain adaptation theory. When tested on a large-scale brain decoding dataset (n=105 subjects), our new mixture model weighting framework achieved state-of-the-art performance—increasing accuracy up to 2.5% over baseline fine-tuning, double the performance gain compared to previous research in supervised source selection. Notably, these improvements were achieved using significantly less training data (i.e., 62% smaller effective sample sizes), suggesting that our performance gains stem from reduced negative transfer. Collectively, this research advances toward a more principled and generalizable brain decoding framework, laying the mathematical foundation for scalable brain-computer interfaces and other applications in computational neuroscience.

## I. Introduction

**B**RAIN decoding is at the forefront of neuroscience and artificial intelligence, driving applications from research in computational neuroscience, to brain-computer interfaces, and to building next-generation machine learning models. Decoding algorithms are used to predict behaviour, cognitive states, and stimuli from patterns of neural activity in the brain [1]–[3]. While traditional statistical pattern recognition algorithms are still used, deep learning models are gaining traction for their ability to extract more complex, non-linear features and improve accuracy [1], [2]. This shift has been fueled by increased access to high-performance computing and the rise of large-scale, multi-subject neural datasets, creating the need for specialized learning algorithms that can handle the inherent variability and high-dimensionality of brain data.

In many deep learning applications, model performance is improved by training on large, diverse datasets from multiple sources. However, in brain decoding, this approach is hindered by negative transfer, where combining data from different subjects can degrade model performance. This occurs because brain activity is highly subject-specific, making it challenging for a single model to generalize across different subjects [4]–[7]. As a result, most pretrained models cannot be directly applied to new subjects and require retraining on large subject-specific datasets, which is time-consuming and inefficient. This challenge exemplifies the broader problem of **multi-source domain adaptation** [8], where the goal is to use data from multiple source domains to improve performance on a target domain. Accordingly, minimizing the impact of negative transfer via multi-source domain adaptation has significant implications for brain decoding, and thus is the focus of our research.

For domain adaptation in brain decoding, the *target domain* refers to the subject for which we want to optimize performance and the *source domains* are the subjects that provide samples for model training. Negative transfer arises from both the learning algorithm’s ability to identify transferable features and the similarity between domains [8]. To mitigate negative transfer in brain decoding, two main methods have been explored: 1) feature alignment and 2) source selection.

Feature alignment seeks to identify transferable features, assuming that an ideal cross-domain representation is one where features are indistinguishable across domains [9], [10]. This can be achieved by minimizing a divergence measure between features from different domains, either directly through adversarial learning to reduce the model’s ability to classify the domain of origin [11] or indirectly via optimization of a proxy objective [12]. However, making domains indistinguishable may prevent the algorithm from learning target-domain specific features that are important for brain decoding [13].

Source selection, in contrast, addresses domain variability by selecting source domains that are more similar to the target domain, thus homogenizing the training dataset without suppressing domain-specific features. Methods for source selection in brain decoding mainly differ in how they define similarity and select source subjects. For example, [14] used cross-subject accuracy as a heuristic similarity measure, selecting the top *k* source subjects most similar to the target subject. The value of *k* was applied globally to all source sets and determined through hyperparameter optimization, balancing generalization benefits with the risk of negative transfer.

Instead of using a fixed *k*, [15] recently proposed a heuristic similarity measure based on the symmetric Kullback-Leibler (KL) divergence. They selected all source subjects with a KL divergence below a specified threshold, *t*, allowing the number of source subjects to vary per target subject. However, *t* remained globally fixed, which does not fully account for target-specific variations. This method also requires hyperparameter optimization to determine a cut-off threshold and uses binary inclusion/exclusion heuristics for source selection rather than continuous weighting.

Unlike the aforementioned studies, [16] selected only the most similar source subject for each target subject, effectively setting *k* = 1 without requiring hyperparameter optimization. Although computationally efficient, this method is limiting, as both [14] and [15] found that the optimal number of source subjects is greater than one. Overall, the prevailing methods for source selection in brain decoding have largely been constrained by their reliance on binary inclusion/exclusion of source subjects and determining the optimal number of source subjects.

Motivated by these limitations, here we reframe source selection for domain adaptation in brain decoding as a parameter estimation problem within a statistical mixture model (Figure 1), where each source subject contributes through a continuous mixture weight rather than being simply included or excluded. This eliminates the need for cut-off thresholds and hyperparameter optimization, uniquely allowing all source subjects to contribute based on their calculated similarity. To compute these weights, we developed a novel convex optimization algorithm based on the Generalized Method of Moments. By using model performance metrics as the generalized moment functions in our optimization, our algorithm also more closely aligns with the theoretical foundations of domain adaptation, enhancing optimality guarantees. Taken together, this research advances toward a more principled and generalizable brain decoding framework, laying the mathematical groundwork for scalable brain-computer interfaces and other applications in computational neuroscience.

**Fig. 1.**
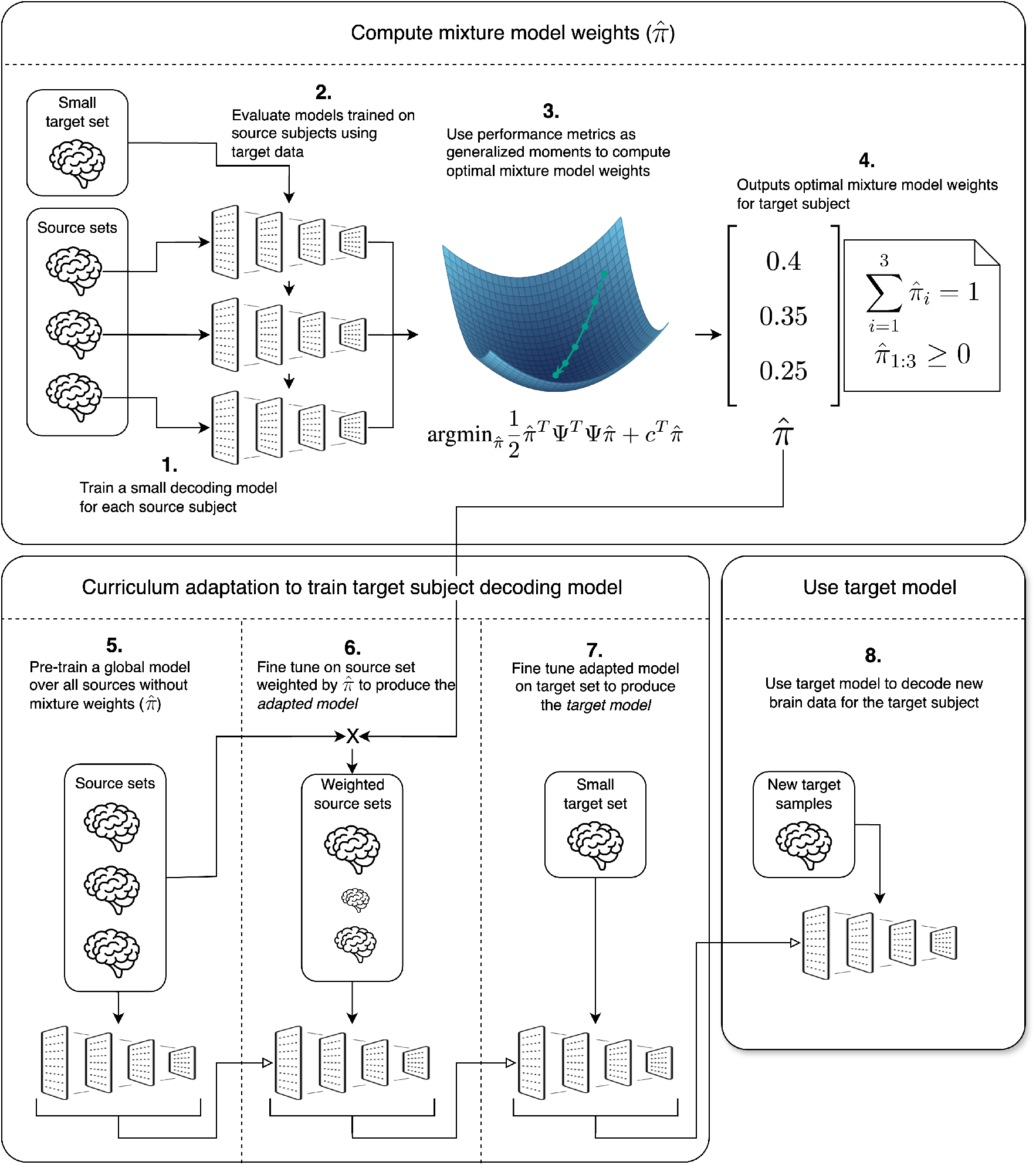
System diagram of our new mixture model weighting framework used for domain-adaptive brain decoding. In phase 1 (top), our source-specific models (1) are evaluated on target subject data (2) to generate generalized moments. Our GMM optimization (3) uses these moments to compute the optimal mixture model weights 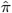 (4). In phase 2 (bottom), a global model is trained over the entire source set without weighting (5), then fine-tuned over the source set weighted by 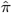 (6), and further fine-tuned on the target data (7) before being used to decode new brain activity for the target subject (8).

## II. Methods

### A. Dataset and Neural Network

We used the large-scale brain decoding dataset by [17] to develop and test our framework. The full dataset includes brain data from 109 subjects, which we reduced to 105, excluding four subjects with variable sample counts. The dataset was recorded using the BCI2000 system with 64 channels and a sampling rate of 160Hz. We extracted 3-second non-overlapping windows, resulting in 84 samples per subject (i.e., 21 samples per class). We used a stratified 80/20 train-test split followed by 5-fold cross-validation, which produced 55 training samples, 13 validation samples, and 16 test samples for each subject. For task classification, we used EEGNet [18], a state-of-the-art neural network model for brain decoding, which uses depth-wise and separable convolutions to minimize parameter count while maintaining competitive performance.

### B. Mixture Model Weights

As the mathematical foundation of our framework, in this section, we show that a target data distribution can be approximated by a weighted mixture of the source data distributions. We show that importance sampling, which underlies domain adaptation theory, can be simplified into a mixture model. We used this model to 1) develop a unifying theoretical framework that positions all source selection methods as special cases, 2) demonstrate that continuous mixture parameters can serve as source domain weights, and 3) develop a computationally efficient algorithm to estimate these weights, using an approach we derived from importance sampling. In machine learning, the main goal is to find a model, *h*(*x*), that minimizes the empirical risk of a loss function, *l*(*h*(*x*), *y*), with respect to a target data distribution, *P*_*T*_ (*x, y*).

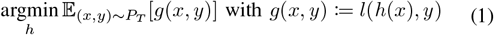

When the target distribution, *P*_*T*_ (*x, y*), is limited or unavailable, importance sampling can provide a principled way to reweight samples from a different distribution, *P*_*S*_(*x, y*), so that optimization can still be performed with respect to *P*_*T*_ (*x, y*). This forms the theoretical foundation of domain adaptation, where models are trained using data from source domains to perform on a target domain [19].

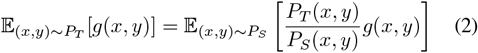

Multi-subject brain decoding is an example of multi-source domain adaptation. Here, the source distribution, *P*_*S*_, is divided into domain-specific distributions, 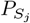, where *j* belongs to the set of source domains *J*. In practice, we do not observe the true distributions *P*_*T*_ and *P*_*S*_, but only finite samples, (*x, y*). Accordingly, these distributions are approximated.

We can approximate the relationship between a source and target distribution using a single, constant importance weight, *π*_*j*_, for each source domain *j*. These constant importance weights follow from systemic similarities and differences between subjects. While real-world distributional shifts are sample-dependent, this first-order approximation allows us to reframe source selection as a parameter estimation problem. Assuming that for each source domain there is a single importance ratio approximation, *π*_*j*_, the expected value of *g*(*x, y*) under the target distribution simplifies to a weighted sum of expectations over the source domains.

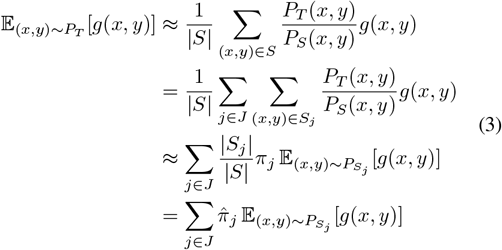

where 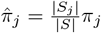 represents a normalized importance weight for domain *j*. As the sample size becomes arbitrarily large, we find

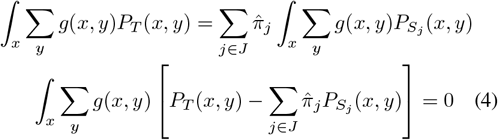

Since this equality holds for any arbitrary function *g*(*x, y*), the underlying distributions must also be equal:

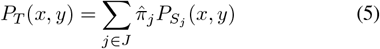

Having a constant importance ratio in each source domain results in an expression in which the target distribution is represented as a mixture model of the source distributions (Figure 2). Traditional binary source selection can thus be seen as a special constrained case where the weights, 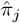, are restricted to be uniform for a subset of subjects and zero otherwise.

**Fig. 2.**
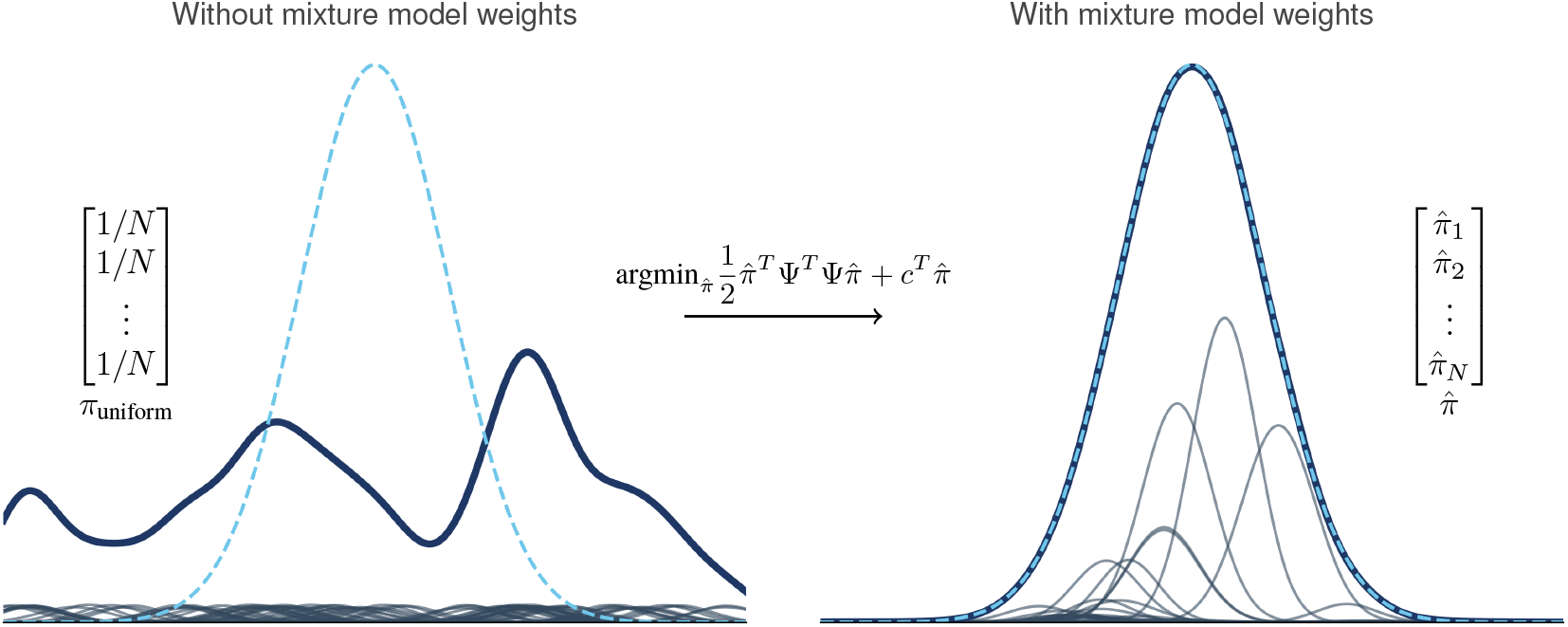
Illustration of data distribution matching using Gaussians with randomized parameters. The standard approach to training a brain decoding model (left) results in uniformly weighted source distributions (grey) that combine into a poorly matched aggregate distribution (blue) relative to the target (dashed). In contrast, our mixture model weighted combination (right) more closely approximates the target data distribution.

Having shown that the target distribution can be mathematically expressed as a weighted mixture of the source distributions (Eq. 5), we then estimate the mixture model weights, 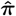. The goal is to choose weights that best approximate the importance sampling ratios defined in Equation 2.

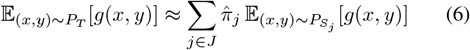

Since *g*(*x, y*) is arbitrary, this expression should hold for any set of functions {*g*_*m*_(*x, y*)}, enabling the use of the Generalized Method of Moments (GMM) to estimate 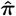. GMM uses an optimization objective to estimate the parameters of a statistical model when *generalized moments*—expectations of functions over the underlying distributions—are available. Using these generalized moments, denoted ℒ_*g,P*_, we rewrite Equation 3:

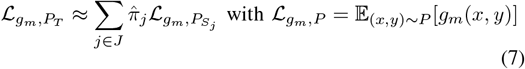

Using the set of arbitrary functions, *g*_*m*_(*x, y*), GMM estimates the weights by minimizing the square error between the generalized moment functions of the weighted source and target data distributions.

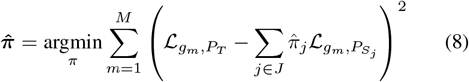

To efficiently solve for the global optimum, we rewrite the moment-matching objective in matrix form (Figure 3), where Ψ ∈ 𝔼^*M ×J*^ is the moment matrix containing the generalized moments of each source distribution and Ψ^Target^ ∈ 𝔼^*M*^ is the vector containing the target generalized moments. This leads to a quadratic optimization problem:

**Fig. 3.**
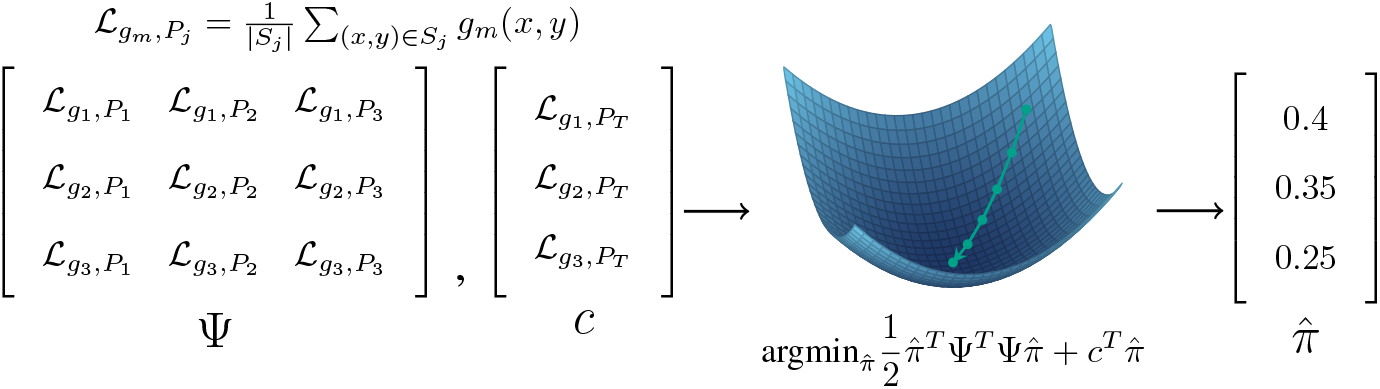
Our new framework for computing source domain importance weights. We used generalized moments,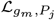, to construct the source moment matrix Ψ and target moment vector *c* for source domains. The optimization objective yields a convex surface with a unique global minimum, producing optimal mixing weights *π* that approximate the target distribution as a weighted combination of the source distributions.

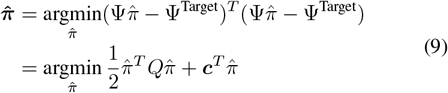

where *Q* = 2Ψ^*T*^ Ψ and ***c*** = − 2Ψ^*T*^ Ψ^Target^. We also constrained 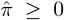 and 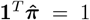 to ensure 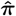 is a valid probability distribution. The globally optimal weights can be computed using quadratic programming with linear constraints. The resulting weights provide a principled and interpretable way to softly include or exclude source domains based on their calculated similarity to the target domain.

Now that we developed an algorithm to estimate the mixture model weights by optimally approximating the importance ratios between the target domain and the set of source domains, we then define the set of functions used to compute the generalized moments, {*g*_*m*_(*x, y*)}. To ensure that the mixture model weights best approximate the importance ratios relevant to domain adaptation, we used the generalized moments from the risk minimization objective in Equation 1.

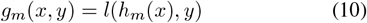

Each *h*_*m*_(*x*) is a brain decoding model trained on a single source subject as illustrated in step 1 of Figure 1 and *l*( ·, ·) is any evaluation metric. When *l*( ·, ·) is a loss function, the expected loss over the weighted source set is approximately equal to the expected loss over the target set. This makes the weighted source and target data distributions approximately equivalent for the purposes of training a brain decoding model.

This led to a principled choice of cross-entropy for *l*, since the loss function is minimized during training. We also evaluated accuracy, which has been used in source selection research for brain decoding [14] and provides robustness to outliers due to its bounded range. For each of the 105 subjects, we treated the other 104 subjects as sources and computed the optimal weights using our mixture model weighting framework. We repeated this process with cross-entropy loss and accuracy as the evaluation metric, resulting in 210 mixture models each comprising 104 source distributions.

### C. Comparative Benchmark

As a comparative benchmark, we implemented the source selection method for brain decoding developed by [14]. Their method ranks potential source domains based on cross-domain accuracy and selects the top *k* performing sources. As we previously showed, the source sets produced by such methods are a constrained case of our mixture model weighting, where all included subjects have the same weight.

To implement their method, we calculated the optimal number of source subjects, *k*, through hyperparameter optimization. For each value of *k*, we constructed a source set for every target subject by selecting the *k* source domains with the highest cross-domain accuracy. We then trained a decoding model for each target subject using its corresponding source set and evaluated on target validation data. We calculated the optimal hyperparameter to be *k* = 20, which yielded the highest average validation accuracy across all target subjects.

In addition to accuracy, we also used loss to rank source subjects. We choose the *k* source subjects with the lowest cross-subject loss. Although [14] did not use loss to rank sources, we included this for comparison with our mixture model weighting. Similar to our framework, we applied their source selection algorithm to each target subject, resulting in 210 mixture models, each comprising 104 source distributions. Overall, we tested our framework using both cross-entropy loss and accuracy as the generalized moment functions, as well as [14] using both cross-entropy loss and accuracy as the evaluation metrics, resulting in 2100 mixture models over 4 experimental conditions.

### D. Curriculum Adaptation

Now that we estimated the target mixture model weights (i.e., step 4 in Figure 1), we used curriculum adaptation to mitigate overfitting while preserving the benefits of domain adaptation. We used the Adam optimizer with *β*_1_ = 0.9 and *β*_2_ = 0.99 and varied the learning rates for different phases of the training: 1 × 10^−3^ for pre-training and 5 × 10^−4^ for the mixture model weighted domain adaptation and subject-specific fine-tuning. We used a batch size of 32 and early stopping with a patience of 5 epochs.

We used a three-phase curriculum adaptation to progressively transition from high-bias, large-sample datasets (global) to low-bias, small-sample datasets (subject-specific). We added a training phase between the standard global pretraining and subject-specific fine tuning, where the decoding model was fine-tuned on our mixture model weighted dataset (i.e., steps 5, 6, and 7 in Figure 1).

1. Global pre-training: We trained five global models, one for each cross-validation split, using uniformly weighted samples from all source domains.
2. Mixture model weighted domain adaptation: We finetuned each global model using our new domain-weighted dataset, producing 2100 total adapted models (5 cross-validation splits × 4 experimental conditions × 105 target subjects).
3. Subject-specific fine-tuning: We fine-tuned each adapted model and global model using target subject data, resulting in 2625 final models adapted for the target subject (2100 from weighted domain adaptation + 525 from fine-tuned global models).

We used this curriculum adaptation to retain both the model generalization induced by pre-training on all subjects and the specificity induced by fine-tuning on a target subject. In an ablation study, we found that removing the regularization effect of global pre-training made the model performance more sensitive to dataset size, masking the effectiveness of reducing negative transfer.

### E. Model Evaluation

We compared 5 experimental conditions: (1) baseline fine-tuning, (2) accuracy-based source selection [14], (3) loss-based source selection [14], (4) our loss-based mixture model weighting, and (5) our accuracy-based mixture model weighting. We evaluated the models using a 16-sample test set for each subject and used accuracy as the main performance metric. Paired t-tests were used to assess improvements in decoding accuracy over baseline fine-tuning. We also tracked the effective sample size (ESS) and overfitting gap.

Effective sample size is used to quantify the diversity of a weighted dataset. In deep learning, a higher ESS is generally preferred, as it is analogous to a larger dataset.

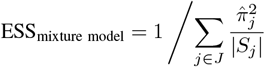

When each data point across all selected domains is weighted equally [14], ESS simplifies to the total number of samples in the selected domains. For fine-tuning on a single domain (i.e., subject-specific tuning), the weight vector becomes a one-hot encoding, where 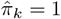 and all other weights are zero.

Overfitting gap is used to quantify model generalization by measuring the difference between training and validation loss. In our study, we used the average difference between the train and validation loss over the last five epochs, where training accuracy was stabilized. Our code is publicly available at https://github.com/Veldrovive/mixture-model-domain-adaptation.

## III. Results

Our mixture model weighting framework achieved state-of-the-art performance for large-scale brain decoding (see Table I). Relative to baseline fine-tuning (66.7%), when using loss as the generalized moment, our new framework increased accuracy by 2.1% (68.8%), whereas previous methods [14] increased accuracy by 0.1% (66.8%). Moreover, when using accuracy as the generalized moment, our framework resulted in a 2.5% increase in accuracy (69.2%) over baseline fine-tuning, whereas previous methods resulted in a 1.3% increase (68.0%).

**TABLE I.** Comparison of our mixture model weighting framework with a fine-tuned baseline and previous research in source selection for brain decoding [14] in terms of test accuracy and loss. The best performance is highlighted in bold

| Metric | Fine-Tuning | Mixture Model (Loss) | Mixture Model (Accuracy) | Source Selection (Loss) | Source Selection (Accuracy) |
| --- | --- | --- | --- | --- | --- |
| Test Accuracy | 66.7% | 68.8% | **69.2%** | 66.8% | 68.0% |
| Test Loss | 0.828 | 0.807 | **0.790** | 0.893 | 0.842 |

Figure 4 shows the weight distributions for two target subjects, including the ranks required to achieve 80% cumulative weight (e.g., rank 33 for target subject 1 and rank 20 for target subject 2). These results show that the number of source domains that meaningfully contribute to the target domain significantly vary across subjects. Calculating the 80% rank across all 105 target subjects resulted in a mean rank of 20.4 ± 6.5. These results also show that the shapes of the weight distributions significantly differ across subjects (e.g., target subject 1 displays a gradual decay, whereas target subject 2 displays a distinct cutoff).

**Fig. 4.**
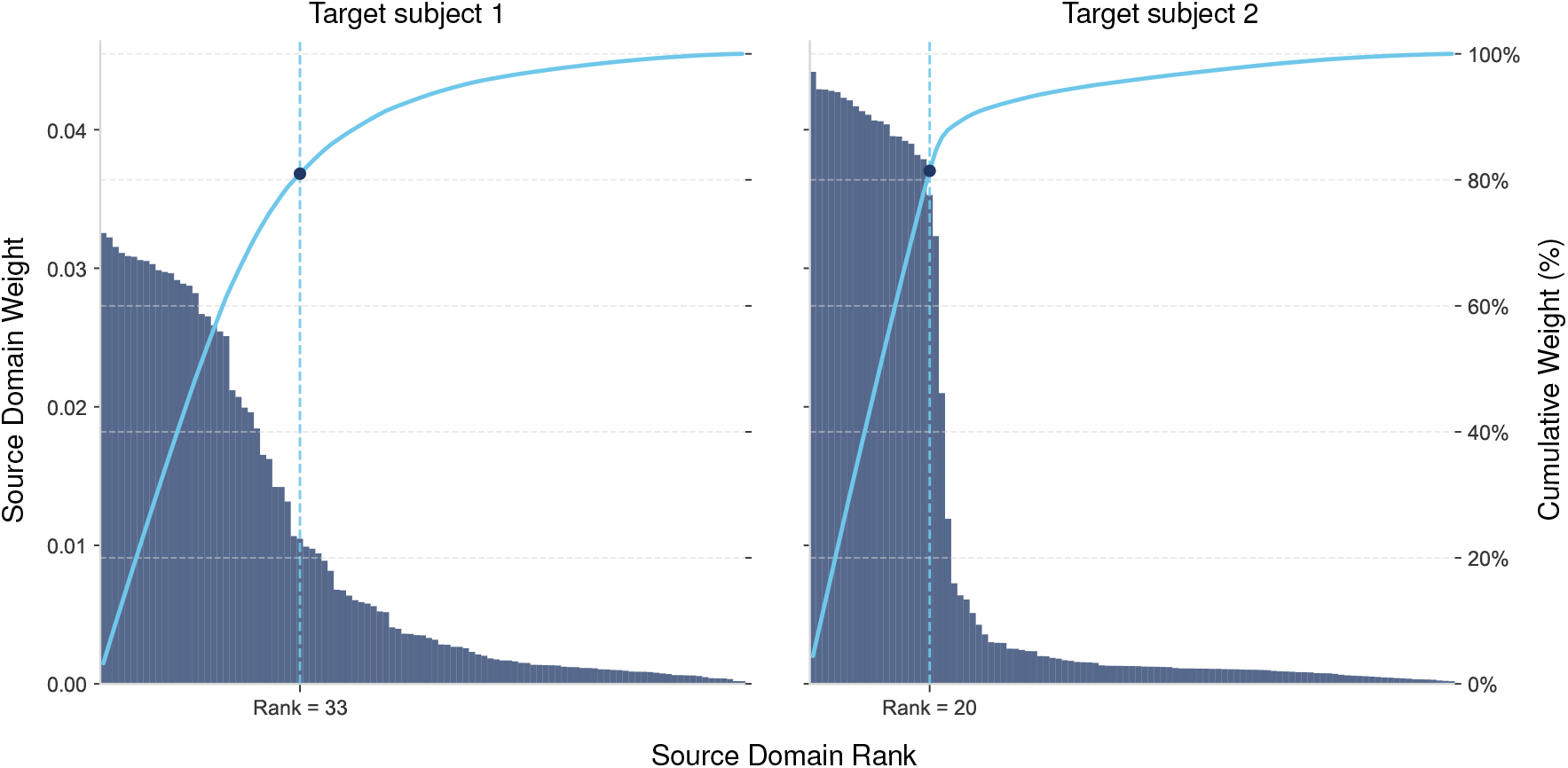
Our mixture model weight distributions for two target subjects. Here, we used accuracy as the generalized moment function. Each bar shows the weight for a single source domain, with domains ranked by weight and plotted in descending order. The curve tracks cumulative weight and the dashed line shows where 80% cumulative weight is achieved.

The results of our curriculum adaptation ablation are summarized in Table II, ordered by descending overfitting gap. Compared to previous source selection research [14], our mixture model resulted in datasets with significantly smaller effective sample sizes (i.e., on average 62% less training data) and greater overfitting, but achieved higher overall test accuracies. This result illustrates that minimizing overfitting alone is not sufficient to maximize generalization performance in large-scale brain decoding.

**TABLE II.** Comparison of our mixture model weighting framework and previous research in source selection for brain decoding [14] in terms of overfitting gap, test accuracy, and effective sample size (ESS)

| Experiment | Pre-Training | Weighting Method | Selection Metric | Overfitting Gap | Test Accuracy (%) | ESS |
| --- | --- | --- | --- | --- | --- | --- |
| 1 | No pre-train | Mixture model | Loss | 0.999 | 59.8 | 377 +/- 158 |
| 2 | Global pre-train | Mixture model | Loss | 0.870 | 68.8 | 377 +/- 158 |
| 3 | No pre-train | Mixture model | Accuracy | 0.744 | 62.4 | 704 +/- 206 |
| 4 | Global pre-train | Mixture model | Accuracy | 0.658 | 69.2 | 704 +/- 206 |
| 5 | No pre-train | Source selection | Accuracy | 0.540 | 60.3 | 1428 +/- 0 |
| 6 | No pre-train | Source selection | Loss | 0.504 | 60.5 | 1428 +/- 0 |
| 7 | Global pre-train | Source selection | Accuracy | 0.444 | 68.0 | 1428 +/- 0 |
| 8 | Global pre-train | Source selection | Loss | 0.437 | 66.8 | 1428 +/- 0 |

## IV. Discussion

In this study, we developed a new computational framework to minimize negative transfer for domain-adaptive brain decoding by reframing source selection as a mixture model parameter estimation problem. Unlike traditional binary inclusion/exclusion heuristics, our framework uniquely allows all source subjects to contribute via continuous weights. We computed these weights using a novel convex optimization algorithm based on the Generalized Method of Moments. When tested on a large-scale brain decoding dataset (n=105 subjects), our framework achieved state-of-the-art performance, increasing accuracy up to 2.5% over baseline fine-tuning, double the performance gain compared to previous research in supervised source selection [14]. Importantly, these gains were achieved using significantly less training data (i.e., 62% smaller effective sample sizes), suggesting that our performance improvements arise from reduced negative transfer. Collectively, this research helps lay the mathematical foundation for developing generalizable brain decoding algorithms, with practical implications for brain-computer interfaces and scientific implications for computational neuroscience.

Our main focus was addressing negative transfer in multi-subject brain decoding. In deep learning, the standard practice to improve model performance is to increase the quantity and diversity of training data. However, due to negative transfer, increasing dataset diversity can sometimes degrade performance, leading to counterintuitive results where models trained on smaller datasets outperform those trained on larger ones [8]. Our results support this principle. Compared to previous research in brain decoding [14], our mixture model weighting framework achieved higher overall test accuracies using training datasets of significantly smaller effective sample sizes.

Although we achieve higher decoding accuracy, our algorithm also exhibits increased overfitting, with an overall validation loss gap of 0.658 (Table II) compared to previous research (0.444). This suggests that our performance may be suboptimal. The lower overfitting in previous research results from hyperparameter optimization that implicitly balances over-fitting and negative transfer by maximizing test performance. In contrast, we focused on minimizing negative transfer without directly addressing overfitting. In both methods, the effective sample size scales with the number of samples per source subject, suggesting that overfitting is a consequence of dataset size. As large-scale neural datasets become more accessible with the rise of commercial and clinical applications [3], [20]–[22], overfitting is expected to decrease, while negative transfer will likely remain a fundamental challenge. Accordingly, the true impact of our framework lies in its ability to tackle this enduring issue, regardless of dataset size.

Our research is a notable departure from previous studies in source selection for brain decoding [14], [16], which assign equal weight to all source domains. As shown in Figure 4 for target subject 2, our mixture model weighting framework distinguishes source subjects into two groups: high-weight and low-weight plateaus, separated by a transition. While previous research approximates this weight distribution by assigning equal weight to all sources on the high-weight plateau and zero to those on the low-weight plateau, we observe significant variability across subjects. For example, target subject 1 shows a gradual transition with intermediate weights for several source domains. This illustrates that binary inclusion/exclusion heuristics may be too rigid, while the adaptability of our framework more accurately captures these subject similarity distributions.

Our research also challenges the traditional assumption of a fixed threshold for inclusion/exclusion, such as that by [14], which uses a single *k* value for all target subjects. This assumes that the same amount of source data benefits all individuals. In contrast, our results show that the optimal number of sources is subject-specific and can vary considerably, with a mean rank of 20.4 ± 6.5 for 80% cumulative weight (Figure 4). While a global *k* = 20 may be optimal on average, it is suboptimal for any individual who deviates from this mean. Our new framework, which uses continuous weights, removes this need for a fixed cut-off threshold, as the mixture model automatically assigns near-zero weights to subjects that would otherwise be excluded.

Despite these advantages, several challenges remain. A critical aspect of our brain decoding framework is its reliance on globally defined source domains. Our theoretical model (Eq. 5) assigns a single importance weight 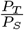 per source domain *j*, enabling tractable optimization and robust moment estimation given typical per-subject dataset sizes. However, this subject-level granularity may overlook finer structures. As noted in subdomain adaptation literature [23], aligning entire domains can obscure discriminative features by merging distinct patterns. A promising extension of our research is applying the mixture model to class-level subdomains, which recent studies suggest can improve performance [15], [23]. Unlike adversarial subdomain alignment that requires end-to-end machine learning, class-level subdomains can be defined pre-training and thus easily compatible with our framework. A challenge, however, will be managing the increased data demands, as finer-grained domains can reduce moment stability and introduce noise in weight estimation.

Lastly, we intentionally isolated source selection from other domain adaptation methods to provide a clean benchmark and detailed performance analysis. While many state-of-the-art systems combine source selection with feature alignment, doing so introduces confounding factors that make it difficult to assess their individual contributions [14], [15]. In practice, these methods are complementary, and combining them is desirable for brain decoding. Future research should consider studying our mixture model weighting in combination with feature alignment. Although we significantly outperformed previous source selection research without feature alignment, it remains to be determined whether this trend holds when other such domain adaptation methods are included.

## V. Conclusion

In conclusion, we developed a new computational framework to address the problem of negative transfer in multi-subject brain decoding. We reframe source selection for domain adaptation as a mixture model parameter estimation problem, allowing each source subject to contribute through a continuous mixture weight. We computed these weights using a novel convex optimization algorithm based on the Generalized Method of Moments, which, using our generalized moment functions, also more closely aligns with the theoretical foundations of domain adaptation, enhancing optimality guarantees. Our framework achieved state-of-the-art accuracy on a large-scale brain decoding dataset using significantly less training data, suggesting that our performance improvements stem from reduced negative transfer. Overall, our new mixture model weighting framework helps lay the mathematical foundation for developing generalizable brain decoding algorithms, with applications ranging from brain-computer interfaces to research in computational neuroscience.

## VI. Acknowledgment

This research is dedicated to the students and researchers in Ukraine who have been affected by the ongoing invasion. Their resilience and unwavering commitment to education and learning continue to serve as a beacon of hope and inspiration to the global academic community.

## References

[1] J. I. Glaser, A. S. Benjamin, R. H. Chowdhury, M. G. Perich, L. E. Miller, and K. P. Kording, “Machine learning for neural decoding,” eneuro, vol. 7, no. 4, pp. ENEURO.0506–19.2020, Jul. 2020.

[2] D. Sussillo et al., “A recurrent neural network for closed-loop intracortical brain–machine interface decoders,” Journal of Neural Engineering, vol. 9, no. 2, p. 026027, Apr. 2012.

[3] J. Ye et al., “A generalist intracortical motor decoder,” Feb. 2025.

[4] N. S. Card et al., “An accurate and rapidly calibrating speech neuroprosthesis,” New England Journal of Medicine, vol. 391, no. 7, pp. 609–618, Aug. 2024.

[5] D. Sussillo, S. D. Stavisky, J. C. Kao, S. I. Ryu, and K. V. Shenoy, “Making brain–machine interfaces robust to future neural variability,” Nature Communications, vol. 7, no. 1, p. 13749, Dec. 2016.

[6] D. M. Brandman et al., “Rapid calibration of an intracortical brain–computer interface for people with tetraplegia,” Journal of Neural Engineering, vol. 15, no. 2, p. 026007, Apr. 2018.

[7] O. Shevchenko, S. Yeremeieva, and B. Laschowski, “Comparative analysis of neural decoding algorithms for brain-machine interfaces,” in 2025 International Conference On Rehabilitation Robotics (ICORR), 2025, pp. 222–227.

[8] F. Zhuang et al., “A comprehensive survey on transfer learning,” Proceedings of the IEEE, vol. 109, pp. 43–76, 2019.

[9] S. Ben-David, J. Blitzer, K. Crammer, and F. Pereira, “Analysis of representations for domain adaptation,” in Advances in Neural Information Processing Systems, vol. 19. MIT Press, 2006.

[10] S. Ben-David, J. Blitzer, K. Crammer, A. Kulesza, F. Pereira, and J. W. Vaughan, “A theory of learning from different domains,” Machine Learning, vol. 79, no. 1, pp. 151–175, May 2010.

[11] O. Özdenizci, Y. Wang, T. Koike-Akino, and D. Erdoğmuş, “Learning invariant representations from EEG via adversarial inference,” IEEE access, vol. 8, pp. 27 074–27 085, 2020.

[12] E. Jeon, W. Ko, J. S. Yoon, and H.-I. Suk, “Mutual information-driven subject-invariant and class-relevant deep representation learning in BCI,” IEEE Transactions on Neural Networks and Learning Systems, vol. 34, no. 2, pp. 739–749, 2023.

[13] D. Kostas and F. Rudzicz, “Thinker invariance: Enabling deep neural networks for BCI across more people,” Journal of Neural Engineering, vol. 17, no. 5, p. 056008, Oct. 2020.

[14] W. Wei, S. Qiu, X. Ma, D. Li, C. Zhang, and H. He, “A transfer learning framework for RSVP-based brain computer interface,” in 2020 42nd Annual International Conference of the IEEE Engineering in Medicine & Biology Society (EMBC). IEEE, 2020, pp. 2963–2968.

[15] K. Liu et al., “Enhancing EEG-based cross-subject emotion recognition via adaptive source joint domain adaptation,” IEEE Transactions on Affective Computing, pp. 1–13, 2024.

[16] E. Jeon, W. Ko, and H.-I. Suk, “Domain adaptation with source selection for motor-imagery based BCI,” in 2019 7th International Winter Conference on Brain-Computer Interface (BCI), 2019, pp. 1–4.

[17] A. L. Goldberger et al., “PhysioBank, PhysioToolkit, and PhysioNet: Components of a new research resource for complex physiologic signals,” Circulation, vol. 101, no. 23, Jun. 2000.

[18] V. J. Lawhern, A. J. Solon, N. R. Waytowich, S. M. Gordon, C. P. Hung, and B. J. Lance, “EEGNet: A compact convolutional neural network for EEG-based brain–computer interfaces,” Journal of Neural Engineering, vol. 15, no. 5, p. 056013, Jul. 2018.

[19] A. Farahani, S. Voghoei, K. Rasheed, and H. R. Arabnia, “A Brief Review of Domain Adaptation,” 2020.

[20] E. Musk, “An integrated brain-machine interface platform with thousands of channels,” Journal of medical Internet research, vol. 21, no. 10, p. e16194, Oct. 2019.

[21] G. Kurbis, A. Mihailidis, and B. Laschowski, “An EMG foundation model for neural decoding,” Arxiv, 2025.

[22] B. M. Karpowicz et al., “Few-shot algorithms for consistent neural decoding (FALCON) benchmark,” Sep. 2024.

[23] Y. Zhu et al., “Deep subdomain adaptation network for image classification,” IEEE Transactions on Neural Networks and Learning Systems, vol. 32, no. 4, pp. 1713–1722, Apr. 2021.

